# Cholecystokinin-like Peptide Mediates Satiety by Inhibiting Sugar Attraction

**DOI:** 10.1101/2020.12.14.419390

**Authors:** Di Guo, Yi-Jie Zhang, Su Zhang, Jian Li, Chao Guo, Yu-Feng Pan, Chen-Xi Liu, Ya-Long Jia, Chen-Yu Li, Jun-Yu Ma, Dick R. Nässel, Cong-Fen Gao, Shun-Fan Wu

**Author notes:** These authors contributed equally to this work (Di Guo, Yi-Jie Zhang).

## Abstract

Feeding is essential for animal survival and reproduction and is regulated by both internal states and external stimuli. However, little is known about how internal states influence the perception of external sensory cues that regulate feeding behavior. Here, we investigated the neuronal and molecular mechanisms behind nutritional state-mediated regulation of gustatory perception in control of feeding behavior in the brown planthopper and *Drosophila*. We found that feeding increases the expression of the cholecystokinin-like peptide, sulfakinin (SK), and the activity of a set of SK-expressing neurons. Starvation elevates the transcription of the sugar receptor Gr64f and SK negatively regulates the expression of Gr64f in both insects. This Gr64f regulation is by direct action of SK neurons on Gr64f-expressing neurons of the proboscis and proleg tarsi that co-express the SK receptor CCKLR-17D3. Our findings thus demonstrate how nutritional state induces peptide signaling to modulate sweet perception and thereby feeding behavior.

---

Neuronal control of feeding is interesting for at least two reasons: in the human population there is a growing problem with excess food consumption causing obesity and associated severe health problems, and secondly, pest insects consume large amounts of our crops worldwide. Both problems are very costly to society. Therefore, understanding mechanisms behind regulation of food search, feeding and satiety is of great interest. In general, animal behavior is guided by internal states and external stimuli ^1,2,3,4,5^. Hence, behavioral decisions depend on the integration of signals from the internal and external environment by circuits of the brain. For instance, feeding behavior, which is essential for the survival and reproduction of animals, is regulated by the nutritional state of the organism and depends on the efficacy of chemosensory organs in localizing food in the environment ^3, 6,7,8,9,10,11,12^. Thus, internal nutrient sensors monitor the energy homeostasis and signal hunger or satiety to the nervous system. Hunger signals received by brain circuits are relayed to sense organs to increase their sensitivity. Hence, in a hungry animal the sensory threshold is lowered in olfactory and gustatory receptors that respond to food cues and thereby increases appetitive behavior and food seeking ^6, 10,11,12,13^. Concomitantly, the hunger signals increase the detection threshold for aversive stimuli, such as bitter tastants ^11, 14^. After food ingestion, satiety signals lower the attractive sensory thresholds and also act on neuronal circuits that regulate feeding behavior, thereby stopping further food intake ^12, 15^.

In *Drosophila* and other insects, the modulation of sensory gain in appetitive behavior is largely dependent on neuropeptides and peptide hormones ^4, 12, 15,16,17^. These signaling systems also orchestrate animal behavior and link internal and external sensory cues [see ^17,18,19^]. Thus, several neuropeptides are known to trigger appetitive behavior, foraging, and mobilize energy stores, at the same time as they suppress other conflicting behaviors such as sleep and reproductive behavior [see ^17,18,19,20^]. At the sensory level neuropeptides such as short neuropeptide F (sNPF), myoinhibitory peptide (MIP), CCH2amide and tachykinin (TK) regulate the sensitivity of subpopulations of olfactory receptor neurons (ORNs) to promote food seeking in hungry flies ^6, 10, 21,22,23^. Also gustatory neurons in *Drosophila* are modulated by neuropeptides to regulate sugar and bitter sensitivities ^11^. Hence, in hungry flies neuropeptide F (NPF), via dopamine cells, increases sweet sensitivity in Gr5a expressing cells and sNPF decreases bitter sensitivity^11^. NPF and sNPF are also known to regulate aspects of feeding, metabolism and sleep ^24,25,26,27,28,29,30,31^, suggesting action at several levels of the organism. Another neuropeptide known to regulate food intake in *Drosophila* is drosulfakinin, DSK, related to the mammalian peptide cholecystokinin (CCK)^32^. CCK in mammals and sulfakinins (SKs) in insects, signal satiety and decrease food ingestion ^33,34,35,36,37^.

We chose to study SK signaling in regulation of gustatory input, food search and feeding in two insect species, the brown planthopper *Nilaparvata lugens* and the genetic model insect *Drosophila melanogaster*. The brown planthopper is a serious pest on rice in Asia and causes damage costing more than 300 million US dollars annually ^38^. *N. lugens* is a monophagous pest that pierce the rice stem and sucks sap, thereby transferring a virus ^39^ that destroys the plant ^40^. With the magnificent genetic toolbox available, *Drosophila* has been extensively used as a model to study regulation of feeding, metabolism and sensory mechanisms underlying food seeking [see ^4, 12, 13, 15, 17, 41^]. Thus, we combine two insect species in our quest to understand how a neuropeptide regulates gustatory perception of food-related taste and ensuing food search and ingestion.

Using *N. lugens* in an RNA-seq transcriptome screen for altered gene expression after downregulation of SK, we found among others an upregulation of sweet-sensing gustatory receptors and the *takeout (to)* gene. Analysis of manipulations of SK and its receptor SKR, as well as gustatory receptors and *to*, suggest that feeding-induced SK signaling downregulates sweet sensing receptors and that *to* is a mediator of SK signaling in the planthopper. Further experiments in *Drosophila* unravel mechanisms behind the satiety signaling that modulates gustatory neurons. Feeding upregulates *Dsk* transcript and increases spontaneous activity and calcium signaling in DSK expressing MP neurons. Furthermore, feeding downregulates Gr64f expression and starvation increases Gr64f and calcium activity in Gr64f expressing neurons. Optogenetic activation of Gr64f neurons increases the motivation to feed. Besides this, knockdown of *dsk* leads to an upregulation of Gr64f transcript and activation DSK expressing MP neurons inhibit the sensitivity of gustatory neurons. Remarkably, we found expression of the DSK receptor CCKLR-17D3 in a subpopulation of the Gr64f GRNs in proleg tarsi, proboscis and maxillary palps and this receptor is downregulated in the appendages after feeding. Finally, knockdown of 17D3 in sweet sensing GRNs decreases the PER. Thus, in summary DSK signaling modulates the sensitivity of sweet-sensing GRNs in a nutrient-dependent fashion in both the planthopper and *Drosophila*.

## Results

### Sulfakinin (SK) reduces food intake in *N. lugens*

In several insect species, SK is known to inhibit feeding and act as a satiety factor ^32,33,34, 42,43,44,45,46^. To test whether SK induces satiety also in *N. lugens*, we injected 4^th^ instar nymph with 20 pmol of either of four types of NlSKs, namely nsNlSK1 (non-sulfated SK1), sNlSK1 (sulfated SK1), nsNlSK2 (non-sulfated SK2) and sNlSK2 (sulfated SK2), dissolved in PBS. After 24h we compared the food uptake to a control group, which was injected with PBS. *N. lugens* injected with sNlSK1 or sNlSK2 consumed 50%-70% less food than the PBS-injected control. However, non-sulfated SKs (nsNlSK1 and nsNlSK2) had no impact on the feeding behavior of *N. lugens* (Fig.1A).

**Fig 1.**
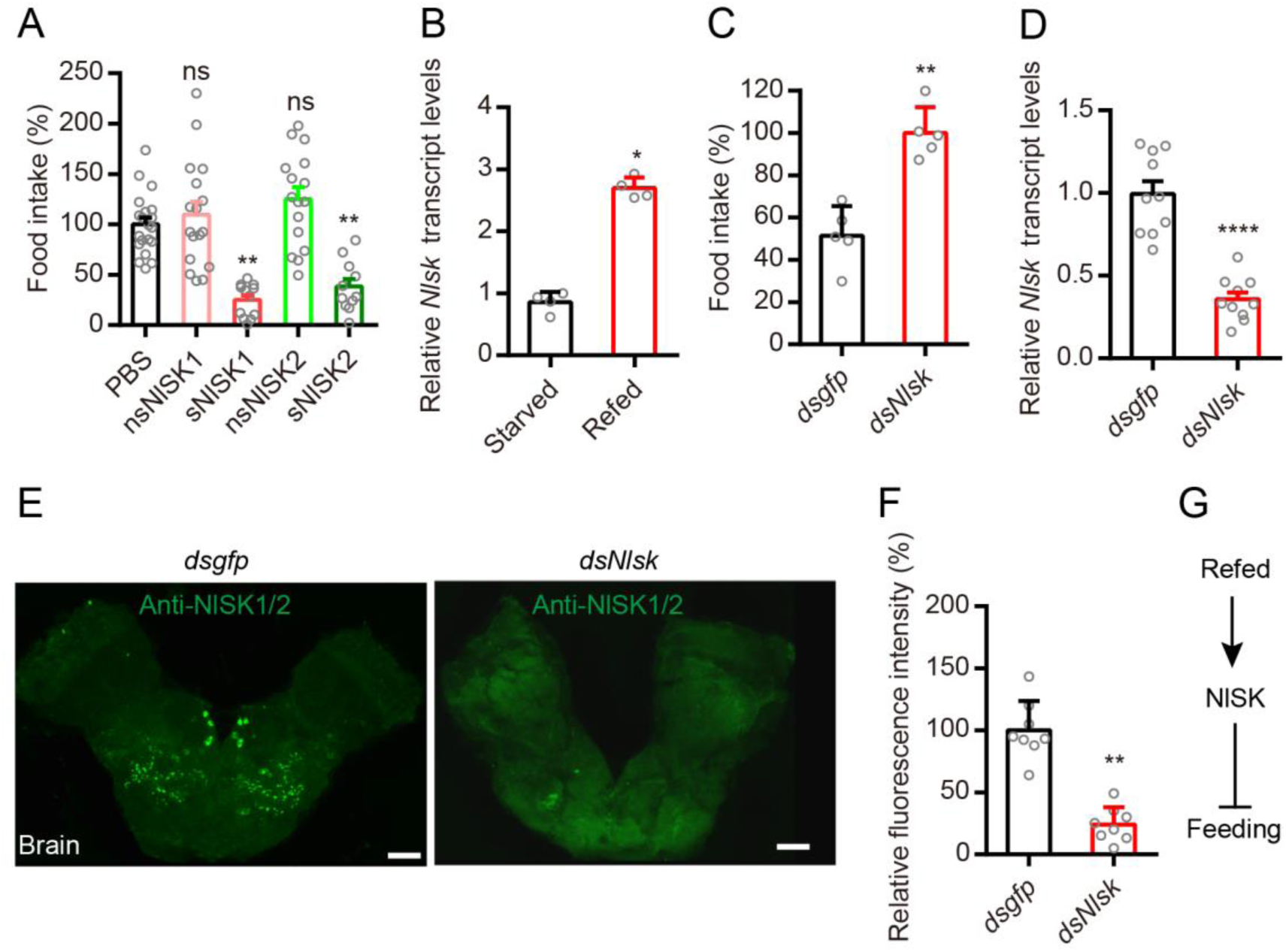
Sulfakinin (NlSK) and its receptor (SKR) signal satiety and inhibit feeding in the rice planthopper. (A) Injection of *N. lugens* sulfated sulfakinins inhibits food intake in the brown planthopper. Fifty nanoliter of PBS (as control) and four sulfakinins (20 pmol/insect) were injected into 4^rd^ instar nymph of brown planthopper. Non-sulfated Sulfakinins (nsNlSK1 and 2) have no effect. All data are presented as means ± s.e.m in this manuscript. ns: not significant; Kruskal–Wallis test followed by Dunn’s multiple comparisons test. (B) Refeeding for 5 hr after 24 hr starvation increases *Nlsk* mRNA transcript. \**p* < 0.05; Mann–Whitney test. (C) Downregulation of *Nlsk* gene using *Nlsk*-RNAi (*dsNlsk*) increases the food intake. \*\**P* < 0.01; Mann–Whitney test. (D) Downregulation of *Nlsk* gene using *Nlsk*-RNAi leads to a reduction in mRNA expression level. \*\*\*\**p* < 0.0001; Mann–Whitney test. (E and F) Downregulation of *Nlsk* gene using *Nlsk*-RNAi (*dsNlsk*) leads to a reduction in NlSK1/2 immunoreactivity. In (E) we show NlSK1/2 immunoreactivity of brain in *dsgfp*- or *dsNlsk*-injected planthoppers. Scale bar: 50 µm. (F) Intensity of anti-NlSK1/2 immunoreactivity in brain. All data are presented as means ± s.e.m. \**p* < 0.05; Mann–Whitney test. (G) Model showing that NlSK inhibits feeding. See also Figure S1-S4

### Effects of *Nlsk* gene silencing on food intake

Next, we asked whether the gene expression level of *Nlsk* is affected by the satiety state. Indeed, we found that the *Nlsk* mRNA level is significantly higher in brown planthoppers that had been refed after 24 h starvation, compared to starved ones (Fig. 1B). To determine the role of *Nlsk* gene in feeding behavior, we performed RNAi injection experiments. Feeding experiments showed that the *dsNlsk*-injected planthoppers consumed approximately 2-fold more food than the *dsgfp*-injected controls (Fig. 1C). The efficacy of the RNAi was tested by qPCR of whole animals and we found that the expression of *Nlsk* was significantly reduced (Fig. 1D). The NlSK1/2 immunoreactivity in the brain was also significantly reduced by the RNAi (Fig. 1E and F). Taken together, these results show that NlSK signaling reflects the nutritional state of the animal and inhibits feeding behavior (Fig. 1G).

### Global gene expression profiling of *N. lugens* in response to *Nlsk*

#### gene-silencing

To unravel the molecular mechanisms underlying control of feeding behavior by *Nlsk* signaling, we performed transcriptome expression profiling (RNA-seq quantification) of *N. lugens* after *Nlsk* knockdown by dsNlsk injection. Illumina sequencing libraries were constructed by using mRNA from the *dsNlsk*- or *dsgfp* (control)-injected 4^th^ instar brown planthoppers. We obtained 49,055,819.5 and 47,254,495 clean reads on average from samples of *Nlsk*–depleted and control groups, respectively (Supplementary Table S2). After removing low-quality regions, adapters, and possible contamination, we obtained more than 6 giga base clean bases with a Q20 percentage over 96%, Q30 percentage over 92%, and a GC percentage between 49.12 and 51.53% (Supplementary Table S2). After alignment by Bowtie, 61.01–65.66% and 61.97–66.21% unique reads were mapped into the reference genome of *N. lugens* (Supplementary Table S3). All of the RNA sequence data in this article have been deposited in the NCBI SRA database and are accessible in PRJNA657327 (SRR12460889-96).

The mapped reads were used to quantify the expression profiling by FPKM (fragments per kilobase of transcript per million mapped fragments) method (Supplementary Table S4). The correlation among four biological replicates in each treatment was then assessed, in terms of the whole genomic FPKM value by using the Pearson method ^47^. The heatmap reveals that the square of correlation value ranged from 0.886 to 0.925 and from 0.861 to 0.916 in the ds*Nlsk*–injected and ds*gfp*-injected control groups, respectively (Fig. 2A). To identify differentially expressed genes (DEGs) in response to *Nlsk* silencing, the DEGs were screened according to the Noiseq method, which is effective in controlling the rate of false positives ^48^. With a cutoff of |Log_2_ fold change| > 1 and *p* < 0.05, a total of 190 DEGs were identified in *Nlsk*–depleted brown planthoppers, including 111 up-regulated and 79 down-regulated DEGs (Fig. 2B and Supplementary Table S5).

**Fig 2.**
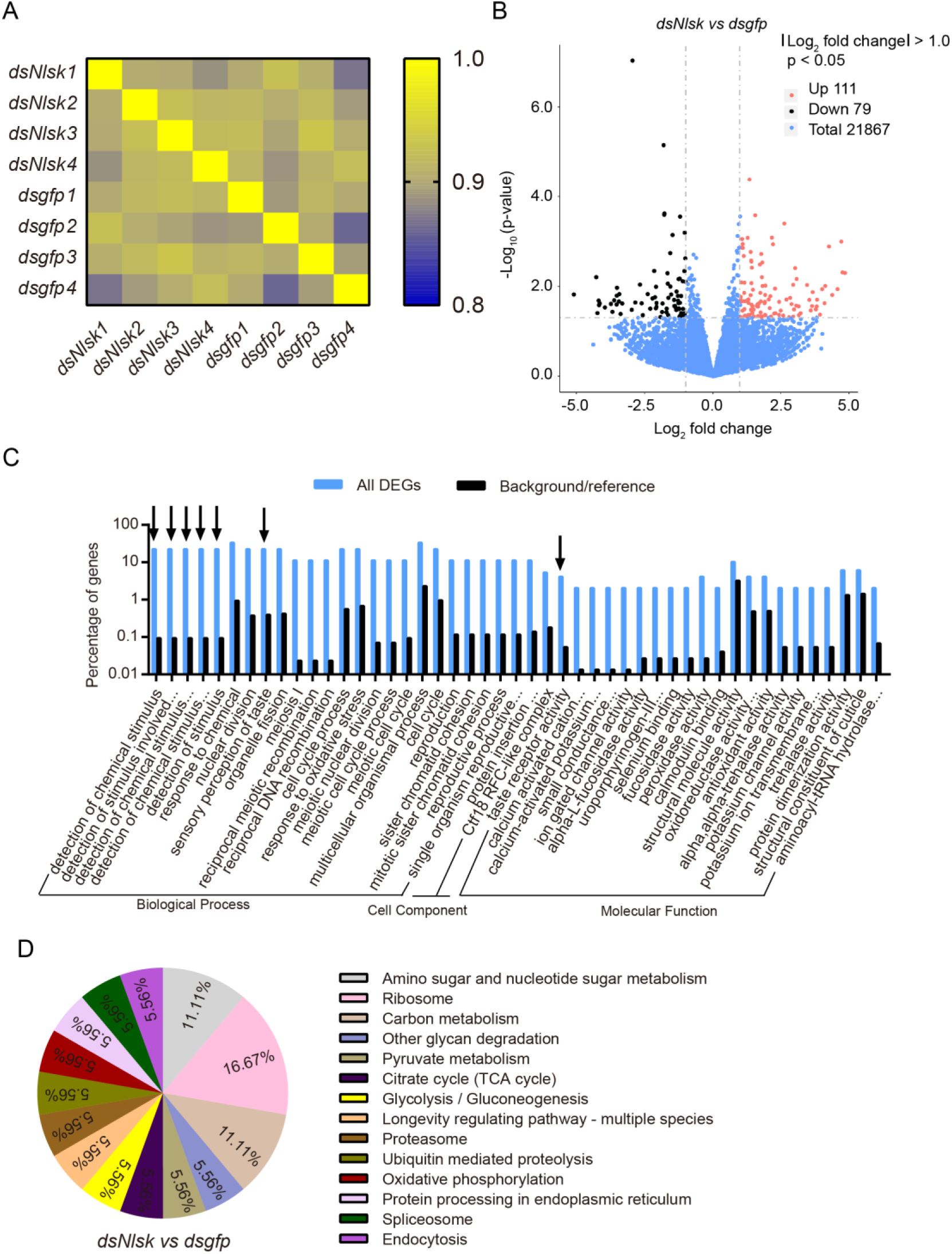
Global gene expression profile in the rice planthopper after *Nlsk* gene knockdown. (A) Heatmap showing the square of correlation value from four biological replicates of *dsNlsk* and *dsgfp* groups analyzed by RNA-seq. The square of correlation value was assessed by using the Pearson correlation. (B) Volcano plot of differentially expressed genes comparing the *dsgfp* and *dsNlsk* treated brown planthoppers. The red dots indicate significantly (*p* < 0.05 and > 2-fold) upregulated genes. The black dots indicate significantly (*p* < 0.05 and > 2-fold) downregulated genes. (C) WEGO (Web Gene Ontology Annotation Plotting) output for *Nlsk* regulated genes. The histogram shows the percent of genes with GO terms enriched in each category. Arrows point to interesting GO categories affected by *Nlsk* knockdown. (D) The pie chart shows the percentage of regulated transcripts distribution in different pathways/processes identified by KEGG pathway analysis.

To obtain a functional classification, these DEGs were assigned to three main GO (Gene Ontology) categories: biological processes (37, 71.42%), cellular components (4, 14.29%), and molecular functions (5, 14.29%) which were further organized into 47 subcategories. (Fig. 2C, Supplementary Table S5). Furthermore, GO analysis revealed an enrichment of GO terms related to detection of chemical stimulus, detection of stimulus involved in sensory perception, detection of chemical stimulus involved in sensory perception of taste, detection of stimulus and taste receptor activity (Fig. 2C) suggesting that genes involved in these processes are regulated by *Nlsk*. For further categorization, the annotated genes were also mapped to KO (KEGG orthologous) terms in the KEGG database. In total, 50 genes (26.32%) could be accessed with a KO number (Supplementary Table S5) and were significantly enriched in several vital physiological processes associated with sugar and carbon metabolism (Fig. 2D). For example, we identified upregulation of a gene encoding the putative juvenile hormone (JH) binding protein *takeout* (*Nlto*, Gene ID: 111048647, 111048649, and 111045963), the ortholog of which, in *Drosophila*, plays an important role in circadian activity and feeding behavior ^49, 50^. Other upregulated genes of interest are associated with sensory perception of sweet taste, including gustatory receptors (Gr) for sugar taste Gr64f-like (NlGr64f, 111045484) and Gr43a-like (NlGr43a, 111056166) (Supplementary Table S4 and S5) ^51,52,53^. Furthermore, genes coding for NADH-cytochrome b5 reductase 2-like (CYB5R2, 111046783), and NADH-cytochrome b5 reductase 4-like (CYB5R4, 111052735), the orthologs of which, in mice, play an important role in sugar metabolism ^54^, were down-regulated. Meanwhile, the *Nlsk* gene (*Nlsk*, 111047610 and 111060391) as positive indicator of our transcriptome data, was significantly repressed after *Nlsk* gene silencing (Supplementary Table S4 and S5). To validate our RNA-seq data, we performed qRT-PCR on six selected genes, three up-regulated and three down-regulated genes in the *dsNlsk*-injection (Fig. S5). The qPCR results are consistent with the RNA-seq data (Supplementary Fig. S1 and Supplementary Table S4).

### *Nlsk* positively regulates *takeout* gene expression that inhibites the feeding behavior of brown planthopper

Takeout (To) has been shown to play an important role in the feeding behavior of *Drosophila* ^49, 50^. To confirm whether *Nlsk* impairs the expression of *takeout* gene (*Nlto*), we performed qRT-PCR to quantify the expression level of *Nlto* gene from the *dsNlsk*-injected, *dsNlskr*-injected and *dsgfp*-injected brown planthopper. Our results show that knockdown of *Nlsk* expression significantly down-regulates *Nlto* expression (Fig. 3A). Besides this, we found that injection of sNlSK2 significantly induces expression of *Nlto* gene (Fig. 3B). These results indicate that NlSK signaling positively regulates the expression of *Nlto* gene (Fig. 3C). Next, we asked whether down-regulation of *Nlto* gene could affect expression of *Nlsk*. After silencing of *Nlto* gene, we did not observe any change in the expression of the *Nlsk* gene (Fig. 3D and E). These results indicate that NlTO has little impact on expression of NlSK (Fig. 3F). Since *to* regulates feeding behavior in *Drosophila* ^49, 50^, we were interested to test its role in the planthopper. Indeed, we observed that brown planthoppers with silenced *Nlto* gene consume more food than the *dsgfp*-injected controls (Fig. 3G). Hence, NlTO inhibits food ingestion (Fig. 3H). Together with previous findings, we propose that feeding induces secretion of NlSK, which promotes release of Takeout and inhibition of feeding behavior in the brown planthopper (Fig. 3I).

**Figure 3.**
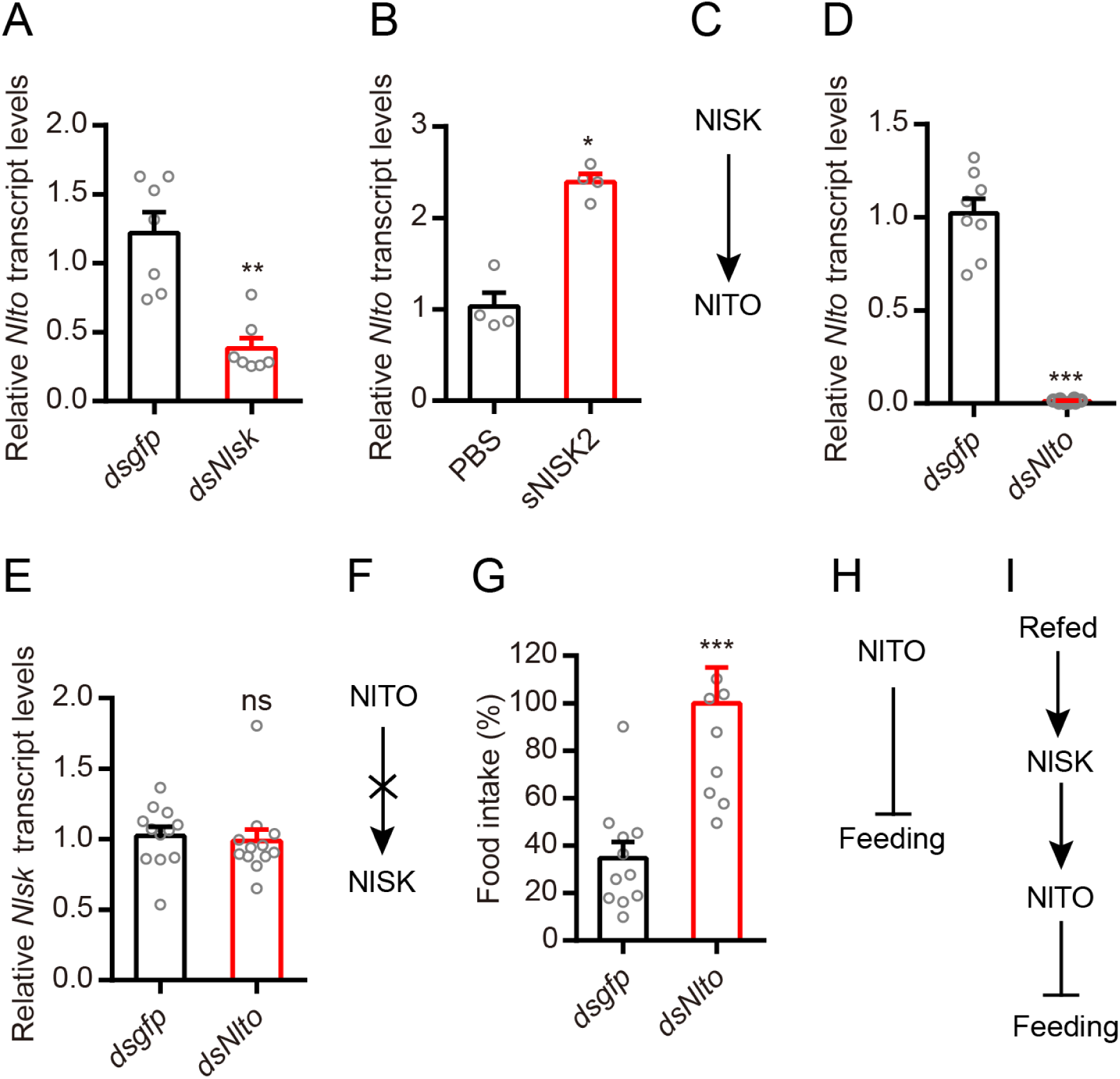
Takeout (to) is a downstream signal of SK that inhibits food intake in the rice planthopper. (A) Knockdown of Nlsk results in downregulation of Nlto gene in the brown planthopper. All data are presented as means ± s.e.m. **P < 0.05; Mann–Whitney test. (B) Injection of sNlSK2 increases gene expression level of Nlto. *P < 0.05; Mann–Whitney test. (C) Model of the NlSK promotes NlTO expression. (D) Downregulation of Nlto gene using Nlto-RNAi leads to a reduction in mRNA expression level. ***P < 0.001; Mann–Whitney test. (E) Knockdown of Nlto have no impacts on expression of Nlsk gene in brown planthopper. All data are presented as means ± s.e.m. ns, no significant difference; Mann–Whitney test. (F) Model showing that NlTO has no effects on NlSK expression. (G) Food intake after silencing Nlto gene. The dsNlto-injected nymphs eat five times more food than dsgfp-injected nymphs in the normal conditions. All data are presented as means ± s.e.m. ***p < 0.001; Mann–Whitney test. (H) Model of the Takeout in the food inhibition. (I) Model of the Takeout as the downstream signal of NlSK involved in the feeding inhibition.

### *Nlsk* negatively regulates gustatory sugar receptor *Gr64f* expression and thereby reduces feeding in the brown planthopper

Interestingly, we found that silencing *Nlsk* gene leads to an up-regulation of gene transcript of the sugar-sensing gustatory receptor *Gr64f* (Fig. 4A). Furthermore, injection of sNlSK2 down-regulates *Gr64f* gene expression (Fig. 4B). These results indicate that NlSK signaling negatively regulates expression of *NlGr64f* (Fig. 4C). Next, we showed that silencing of *Nlto* also increased the expression of *NlGr64f* (Fig. 4D) indicating that NlTO inhibits expression of NlGr64f (Fig. 4E). We found that the expression of *NlGr64f* was also influenced by feeding state and displays a phenotype opposite to that of the *Nlsk* gene. Thus, refeeding after starvation inhibits the expression of *NlGr64f* (Fig. 4F). Then, we asked whether silencing of the *NlGr64f* gene impairs feeding behavior. Down-regulation of the *NlGr64f* gene indeed diminishes food intake compared to control animals (Fig. 4G and H). These results suggest that internal nutritional states influence chemosensory processing in gustatory receptor neurons (GRNs). Starvation negatively regulates the NlSK signaling pathway, which in turn inhibits NlTO expression and promotes *NlGr64f* expression and leads to increased food consumption. Furthermore, refeeding after starvation positively regulates the NlSK and NlTO signaling pathway, which thereby inhibits the expression of NlGr64f and leads to reduced feeding (Fig. 4I). We also found that *Nlsk* manipulations alter the expression of another sweet-sensing gustatory receptor, *NlGr43a*. Injection of *dsNlsk* increased NlGr43a expression, whereas injection of sNlSK2 decreased its expression (Supplementary Fig. S2A and B). However, we focused on Gr64f in the following analysis of *Drosophila*.

**Figure 4.**
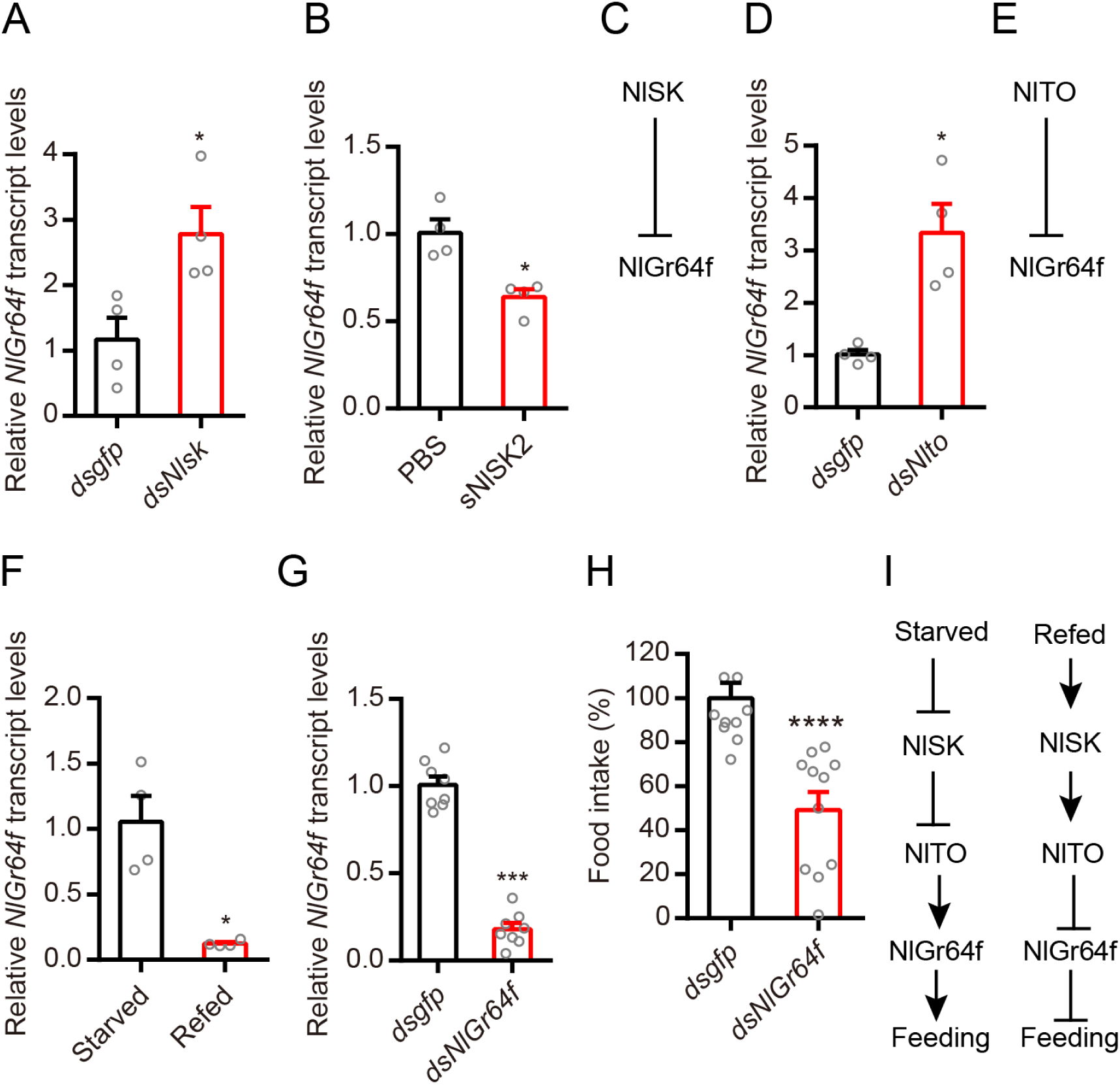
Sulfakinin inhibits expression of the sweet gustatory receptor Gr64f, which promotes food ingestion in the rice planthopper. (A) Downregulation of *Nlsk* gene using *Nlsk*-RNAi (*dsNlsk*) leads to up-regulation of transcript of sweet sensing NlGr64f. \**P* < 0.05; Mann–Whitney test. (B) Injection of sNlSK2 leads to down-regulation of *NlGr64f* gene. \**P* < 0.05; Mann–Whitney test. (C) Model showing that NlSK inhibits NlGr64f expression. (D) Downregulation of *Nlto* gene using *Nlto*-RNAi (*dsNlto*) leads to up-regulation of transcript of sweet sensing NlGr64f. \**P* < 0.05; Mann–Whitney test. (E) Model showing that NlTO inhibits NlGr64f expression. (F) Refeeding for 5 hr after 24 hr starvation decreases *NlGr64f* transcript. \**p* < 0.05; Mann–Whitney test. (G) Downregulation of *NlGr64f* gene using *NlGr64f*-RNAi (dsNlGr64f) leads to a reduction in mRNA expression level. \*\*\**P* < 0.001; Mann–Whitney test. (H) Downregulation of *NlGr64f* gene decreases the food intake. \*\*\*\**P* < 0.0001; Mann–Whitney test. (I) Model showing that the feeding state regulates SK signaling which in turn modulates NlTO and NlGr64f signaling (sweet sensing) and thereby feeding.

### In *Drosophila* Drosulfakinins (DSKs) signal satiety

Next, we used the genetic model insect, *Drosophila melanogaster*, to further investigate mechanisms of DSK signaling in modulation of gustatory sugar reception and feeding. Earlier studies have implicated DSK signaling in regulation of food intake ^32, 36, 55, 56^. First, we found that the *Dsk* mRNA level is significantly higher in flies refed after 24 h starvation, compared to starved flies (Fig. 5A). Consistent with this, immunohistochemistry showed that the level of DSK peptides (anti-DSK1/2) in refed flies is higher than that in starved ones (Fig. 5B and C). To monitor activity in Dsk expressing neurons related to feeding, we next expressed the genetically encoded fluorescent voltage indicator *ArcLight* ^57^ in *Dsk-GAL4* neurons. We observed a higher level of spontaneous activity in the Dsk-expressing MP neurons in refed flies compared to that in starved flies (Fig. 5D and E). Furthermore, we utilised an activity reporter system, calcium-dependent nuclear import of LexA (CaLexA) ^58^, to reveal whether increased activity of the Dsk-expressing MP neurons correlates with refeeding after starvation. Our analysis shows that flies refed after starvation exhibit significantly increased Ca^2+^ activity in the MP neurons, compared to starved flies (Fig. 5F and G). To further confirm our findings, we employed a red-shifted channel rhodopsin, *CsChrimson* ^59^ as an optogenetic effector under control of the Dsk-GAL4. After light activation of Dsk-GAL4 expressing neurons, we found that the PER (proboscis extension response) of flies was significantly decreased compared with control flies (Fig. 5H). We also found that *Dsk* mutants display increased motivation to feed in the capillary feeding (CAFE) assay ^60^ (Fig. 5I). Similar results were also previously reported ^55^. These results suggest that re-feeding after starvation increases DSK expression and activates *Dsk*-expressing MP neurons, which release DSK to inhibit feeding behavior (Fig. 5J).

**Figure 5.**
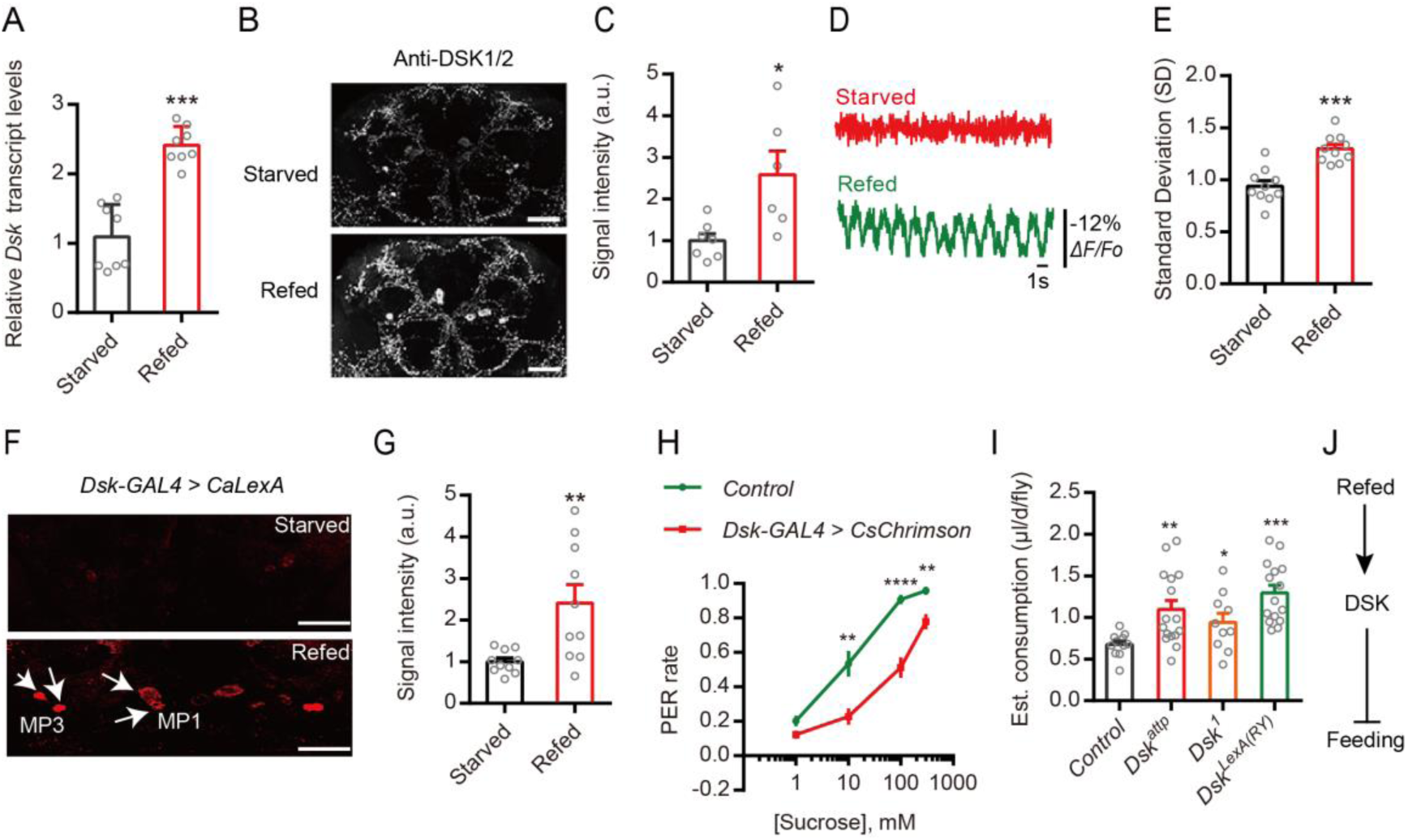
Drosulfakinin (DSK) signals satiety and inhibits feeding in *Drosophila*. (A) Refeeding after 24h starvation increases *Dsk* transcript. \*\*\**p* < 0.001; Mann–Whitney test. (B and C) Refeeding after 24h starvation increases DSK peptide expression as revealed by anti-DSK1/2 immunolabeling. Scale bars, 50 μm. \**p* < 0.05; Mann–Whitney test. (D and E) Refeeding after starvation increases spontaneous neuronal activity of MP1 neurons as monitored by the voltage indicator *ArcLight*. \*\*\**p* < 0.001; Mann–Whitney test. (F and G) Representative images showing CaLexA signals in DSK expressing MP neurons of flies exposed to starvation and refeeding. Starved: flies were raised in 0.5% agar for 24 hours; Refed: flies were raised in fly food 1.5 h after 24 h starvation in 0.5% agar. Red: maximal intensity of *CaLexA* signals. Scale bar, 50 μm. (G) Quantification of the signal intensity of *CaLexA* signals in DSK expressing MP neurons from flies treated with conditions shown in (F). \*\**p* < 0.01; Mann–Whitney test. (H) Optogenetic activation (19 μm/mm^2^) of *dsk-GAL4* neurons expressing *CsCrimson* is sufficient to inhibit proboscis responses. \*\**p* < 0.01, \*\*\*\**p* < 0.0001; Mann–Whitney test. (I) Three Dsk mutants show increased food consumption compared to wildtype in the CAFE essay. \**p*<0.05, \*\**p*<0.01, \*\**p*<0.001, Kruskal–Wallis test followed by Dunn’s multiple comparisons test. (J) Model showing that DSK inhibits feeding.

### The sugar receptor Gr64f promotes motivation to feed in *Drosophila*

We next tested whether sugar-sensing GRNs promote feeding behavior in *Drosophila*. First, we found that the transcript level of *Gr64f* was down-regulated in the refed flies compared with starved ones (Fig. 6A), which is opposite to the *Dsk* transcript (Fig. 5A). Then, we utilized mutants that lacked all eight known sugar receptors (*sugar blind*) ^61, 62^ to perform PER analysis. As expected, sugar blind flies show almost no response to sucrose solutions (Fig. 6B), similar to flies with their sweet GRNs selectively silenced (Fig. 6C). Next, we interrogated whether the activity of sweet neurons is modulated by nutritional state as indicated in former studies ^8, 12^. We selectively expressed the genetically encoded Ca^2+^ indicator GCaMP6s ^63^ in *Gr64f-GAL4* labeled neurons and measured the response to nutrients. The brains of starved or refed flies were imaged for GCaMP signal *ex vivo* (Fig. 6D and E). The GCaMP signal is elevated in the sweet GRNs of starved flies compared to refed controls, suggesting these neurons are more active during starvation (Fig. 6D and E). Next, we tested whether activation of sweet neurons is sufficient to induce feeding behavior. We expressed the optogenetic effector *CsChrimson* under control of the *Gr64f-GAL4*. As with the preceding results, control animals display a low rate of PERs upon stimulation with light stimuli. However, the *Gr64f-GAL4, UAS-CsChrimson* animals show a prominent dose-dependent PER responses under light stimuli (Fig. 6F and G, Supplementary Movie S1). These results indicate that starvation promotes Gr64f expression and activates sugar-sensing GRNs, which reflects a starved state that promotes feeding behavior (Fig. 6H).

**Figure 6.**
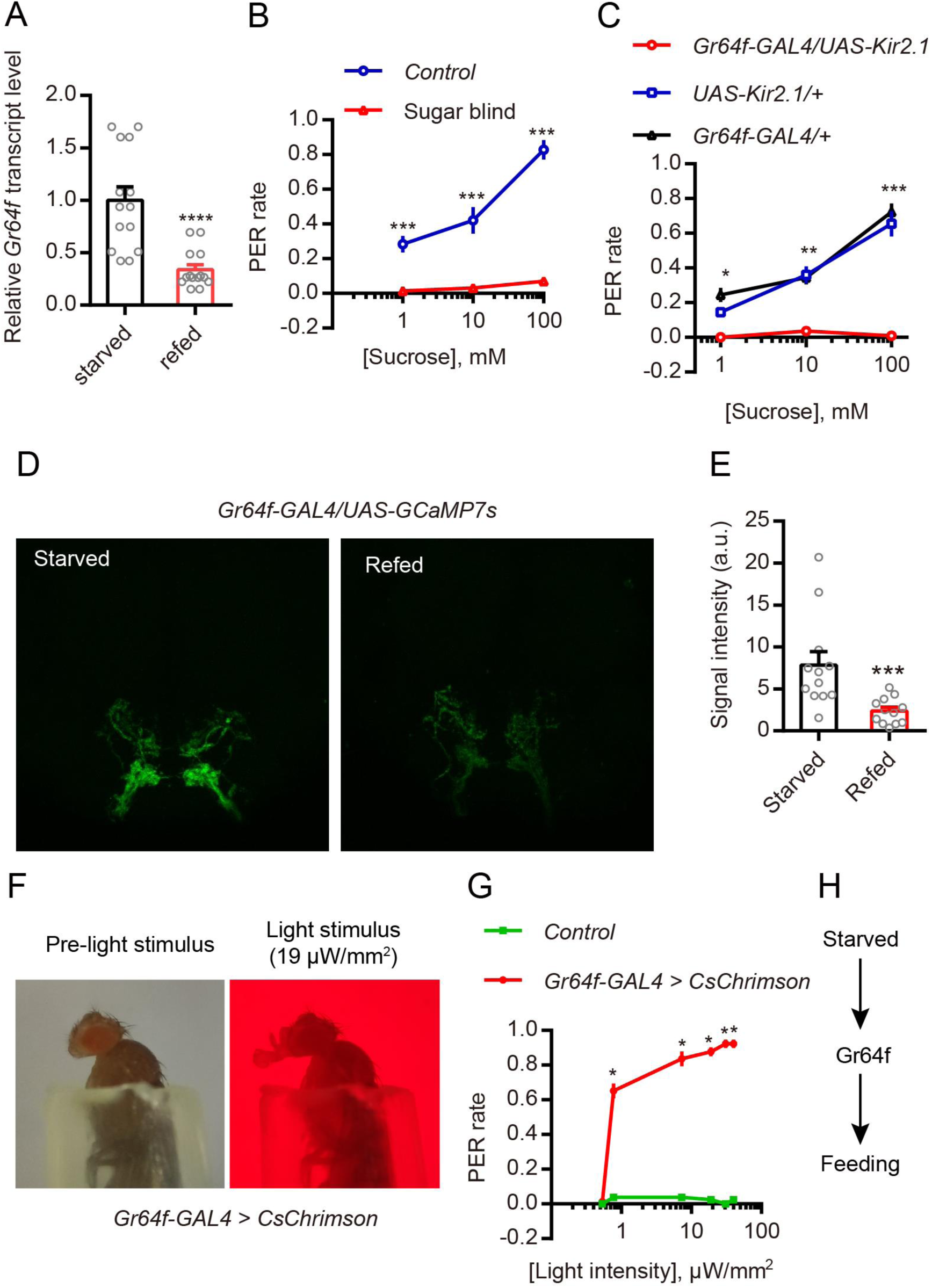
In *Drosophila*, Gr64f promotes feeding. (A) Refeeding after 24h starvation decreases *Gr64f* transcript. \*\*\*\**p* < 0.0001; Mann–Whitney test. (B) The sugar blind mutant (all eight known sugar receptors mutated) show almost no motivation to feed in proboscis extension reflex (PER). \*\*\**p* < 0.001; Mann–Whitney test. (C) Silencing *Gr64f-GAL4* neurons by expressing the hyperpolarizing channel Kir2 also caused loss of motivation to feed in PER. \*\*\**p* < 0.001, \*\**p* < 0.01, \**p* < 0.05; Mann–Whitney test. (D) Representative images of Gr64f-GAL4 drive GCaMP7s fluorescence at SEZ in both starved and refed condition. (E) Quantification of GCaMP7s signal intensity in starved and refed conditions. \*\*\**p* < 0.001; Mann–Whitney test. (F) Dynamics of light-induced proboscis extension after photoactivation in a fly expressing *Gr64f-Gal4*, *UAS-CsChrimson*. (G) Relationship between the intensity of the stimulating light and PER rate using the indicated flies. \**p* < 0.05; Mann–Whitney test. (H) Model showing that Gr64f promotes feeding.

### DSK and TO negatively regulates sugar receptor *Gr64f* expression in *Drosophila*

To test whether silencing *Dsk* gene expression affects the expression of *Gr64f* also in *Drosophila*, we used a UAS-Dsk-RNAi transgene targeted either globally, to Dsk-expressing neurons, or to insulin-producing cells (IPCs) ^64^. Knockdown with the global GAL4 driver (Actin5C-GAL4), and the specific driver (Dsk-GAL4), resulted in up-regulation of the *Gr64f* gene (Fig. 7A and B). However, diminishing of *Dsk* in the insulin producing cells (IPCs) using Dilp2-GAL4 (Supplementary Fig. S3) has no significant effect on *Gr64f* gene expression (Fig. 7C). Our previous studies showed that Dsk-GAL4 is expressed in two sets of MP neurons and a subset of the IPC neurons ^32, 64^, and our data here indicate that DSK from MP neurons, but not from IPCs is responsible for affecting *Gr64f* expression. Similar to brown planthopper, in *Drosophila*, silencing of *Dsk* gene negatively regulates takeout expression (Fig. 7D). Thus, we asked whether *takeout* also negatively regulates Gr64f expression. Indeed, our results show that reducing *takeout* expression using an Actin5C-GAL4 driver upregulates *Gr64f* expression (Fig. 7E).

**Figure 7.**
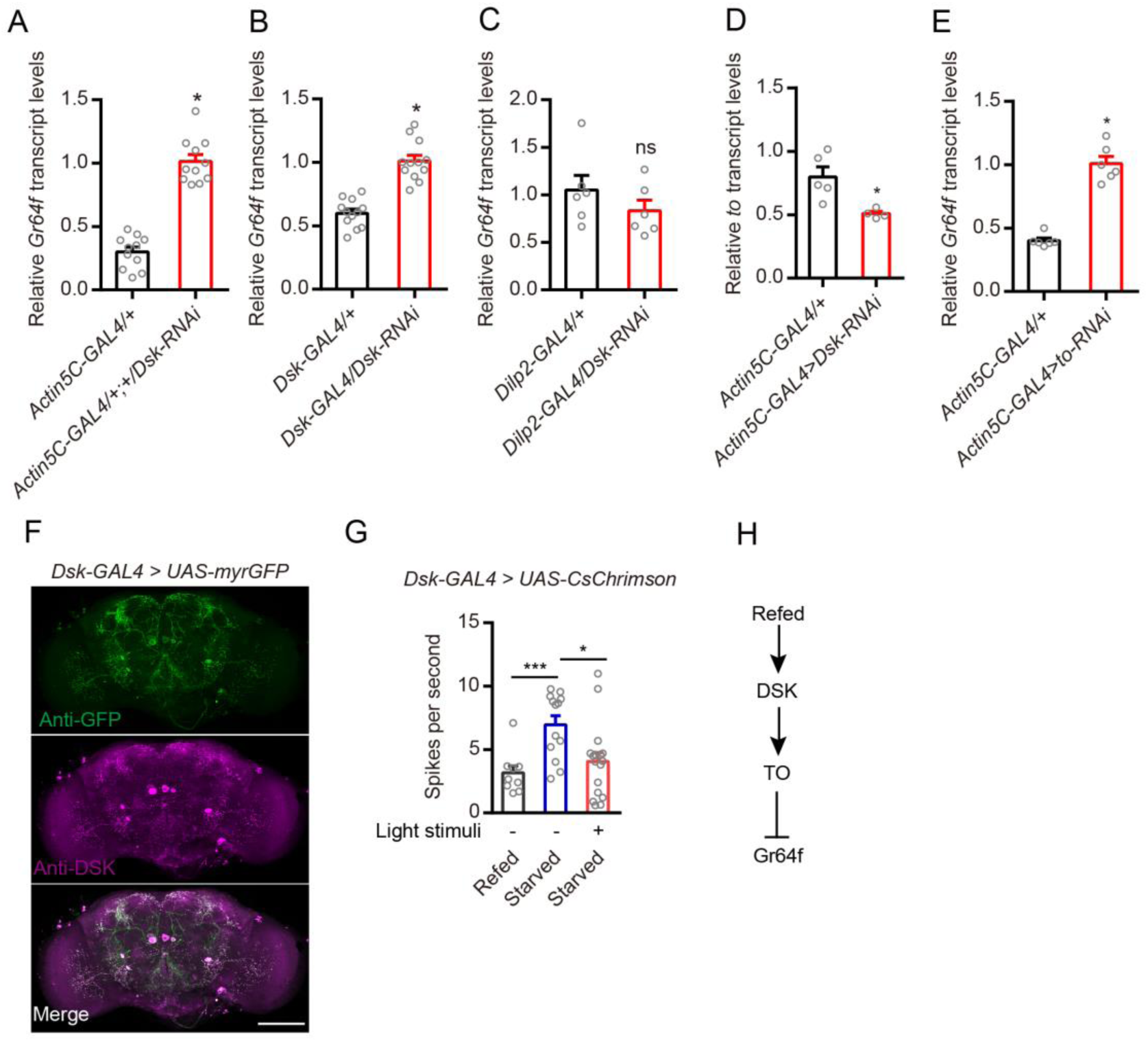
In Drosophila DSK and TO Inhibits Gr64f Gene Transcription and Optogenetic Activation of Dsk-GAL4 Neurons Decreases the Sensitivity of Gustatory Neurons in Starved Flies. (A and B) Relative expression levels of *Gr64f* transcripts in DSK deficient files. Depletion of *Dsk*, either globally in all cells (*Actin5C-GAL4*) or in the Dsk-GAL4-expressing cells increased *Gr64f* transcripts in starved flies. \**P* < 0.05; Mann–Whitney test. (C) Silencing only *Dsk* in the IPCs with RNAi (Dilp2-GAL4/Dsk-RNAi; red bars) did not affect *Gr64f* transcript levels. ns: not significant; Mann–Whitney test. (D) Relative expression levels of *to* transcripts in *Dsk* deficient files. Depletion of *Dsk* in all cells (*Actin5C-GAL4*) decreased *To* transcripts in flies. \**P* < 0.05; Mann–Whitney test. (E) Relative expression levels of *Gr64f* transcripts in TO deficient files. Depletion of *to* in all cells (*Actin5C-GAL4*) increased *Gr64f* transcripts in flies. \**P* < 0.05; Kruskal–Wallis test followed by Dunn’s multiple comparisons test. (F) Expression pattern of *Dsk*-GAL4 in the brain revealed by anti-GFP (upper) and anti-DSK (middle). Scale bars, 50 μm. (G) Electrophysiological recordings reveal that starved flies (24 h) show increased sensitivity of GRNs compared with refed flies (fed 1.5 h after starvation) and optogenetic activation of *Dsk*-GAL4 neurons (30 min) decreases the sensitivity of gustatory neurons in starved flies (24 h). \*\*\**P* < 0.0001; \**P* < 0.05; Mann–Whitney test. (H) Model of the feeding state, Dsk, TO and Gr64f interaction.

### Optogenetic Stimulation of Dsk-GAL4 Neurons Decreases the Sensitivity of Gustatory Neurons in Starved Flies

Next, we set out to identify the potential mechanism by which DSK neurons modulate Gr64f-expressing sugar-sensing GRNs. We found that sugar-sensing GRNs in the proboscis and Dsk-expressing neurons in the brain project their axons to overlapping areas in the subesophageal zone (SEZ) of the brain (Supplementary Fig. S4A and B). We did not detect Dsk-expressing neurons in the proboscis or proleg tarsi (Supplementary Fig. S4C and D). Our Dsk-GAL4 line labels eight MP neurons in the fly brain (Fig. 7F), but, only MP1 neurons project to SEZ of the brain ^64^. Next, we asked whether activation of Dsk-GAL4 neurons could impair the sensitivity of GRNs. As previous studies reported ^50^, we also observed that starved flies show increased sensitivity of GRNs compared with refed flies (Fig. 7G). However, activation of Dsk-GAL4 neurons for 30 min decreases the sensitivity of GRNs in starved flies (Fig. 7G).

These results indicate that refeeding induces release of DSK, which promotes TO and then inhibits sugar receptor expression and sugar-sensing neuron activity (Fig. 7H).

### Distribution of sulfakinin receptor (SKR) in the sweet-sensing gustatory neurons

Earlier studies have shown that GRNs in *Drosophila* are modulated by neuropeptides to adjust sensitivities to sweet and bitter taste ^11^. Hence, in hungry flies NPF, via dopaminergic cells, increases sweet sensitivity and sNPF decreases bitter sensitivity ^11^. Thus, we asked whether DSK receptors are expressed in the sweet-sensing GRNs of the gustatory system.

Two G-protein coupled receptors, the CCK-like receptors (CCKLR)-17D1 and CCKLR-17D3, have been identified as the receptors of DSK peptides in *Drosophila* ^65,66,67^. As we failed to generate functional CCKLR-17D1 and CCKLR-17D3 antibodies, we produced a knock-in GAL4 into the start codon of each of CCKLR-17D1 and CCKLR-17D3 ^64^ for expression analysis (Supplementary Fig. S5A). We did not detect expression of *17D1^GAL4^* in cells of the leg tarsi, maxillary palps or proboscis (Supplementary Fig. S5B). However, we found that 17D3^GAL4^ drives expression broadly in cells of the proleg tarsi, proboscis and maxillary palps (Supplementary Fig. S6). Next, we asked whether the CCKLR-17D3 receptor is co-expressed with sugar receptors in the gustatory receptor neurons (GRNs). Double labeling of *Gr64f^LexA^* and *17D3^GAL4^* expression (*17D3^GAL4^; LexAop-RedStinger, UAS-stinger-GFP/+; +/Gr64f^LexA^*) revealed that five pairs of neurons of the labellum and eight of each proleg tarsi are co-labeled by *17D3^GAL4^* and *Gr64f^LexA^* (Fig. 8A). Thus, we have evidence that CCKLR-17D3, one of the two sulfakinin receptors, is expressed in some of the Gr64f-expressing neurons in the proboscis and leg tarsi (Fig. 8A). We also show here that *Gr64f^LexA^* and *17D3^GAL4^* labeled neurons of the proboscis extend their axons to overlapping areas in the SEZ of the brain (Fig. 8B1 and B2). In addition, we also detected their axons in overlapping areas of the prothoracic neuromere of the ventral nerve cord (Fig. 8B3-5).

**Figure 8.**
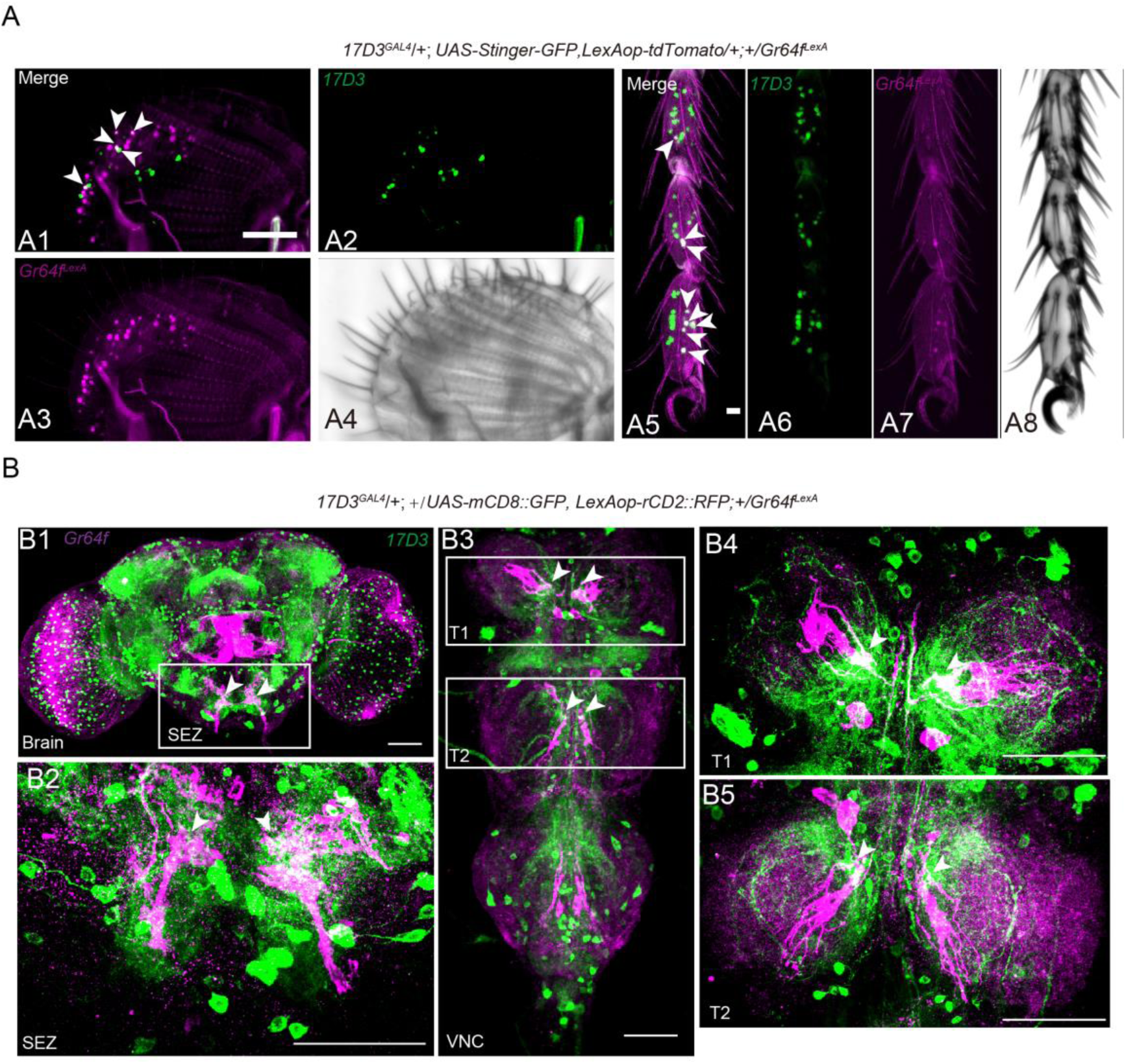
In Drosophila the DSK receptor, CCKLR-17D3, is expressed in Gr64f^LexA^ expressing neurons. (A) *17D3^GAL4^, UAS-Stinger-GFP* (green) superimposes with *Gr64f^LexA^, LexAop > tdTomato* (magenta) expressing cells in the labellum (A1-A4) and proleg tarsi (A5-A8) of flies. Overlap is in white. The white arrowheads (A1 and A5) indicate the positive neural cells co-labeled by *17D3^GAL4^* and *Gr64^LexA^*. Scale bar: 50 μm. (B) Double labeling of 17D3^GAL4^-expressing neurons and Gr64f^LexA^-expressing sweet neurons in the brain (B1), SEZ (B2), and VNC (B3-B5). SEZ: subesophageal zone; VNC: ventral nerve cord; T1-T3: thoracic ganglion 1-3. Scale bar: 50 μm.

### CCKLR-17D3 signaling in GRNs mediates food induced modulation of PER

Next, we asked whether the inhibition of the PER by DSK that we observed is mediated by its receptor, CCLR-17D3, which is expressed by the sweet-sensing GRNs. To address this, we first found that the transcript level of *ccklr-17d3* in the labellum and proleg is down-regulated in the refed flies compared to starved flies (Fig. 9A and B), which is similar to the transcript of Gr64f (Fig. 6A). Secondly, we found that 17D3 mutant flies exhibit decreased PER compared to control flies (Fig. 9C). In contrast, 17D1 mutants do not display a significant difference to controls in our PER assay (Supplementary Fig. S5C). Thirdly, we knocked down either of the DSK receptors (CCKLR-17D1 and CCKLR-17D3) by crossing each of *UAS-17D1-RNAi* and *UAS-17D3-RNAi* with *Gr64f-GAL4*. We measured the PER in our behavioral assay and found that flies lacking 17D3 in sweet-sensing GRNs exhibit a significantly decreased PER, whereas 17D1 knockdown had no effect (Fig. 9D). Thus, 17D3 expression in Gr64f expressing GRNs mediates the starvation-dependent enhancement of PER.

**Figure 9.**
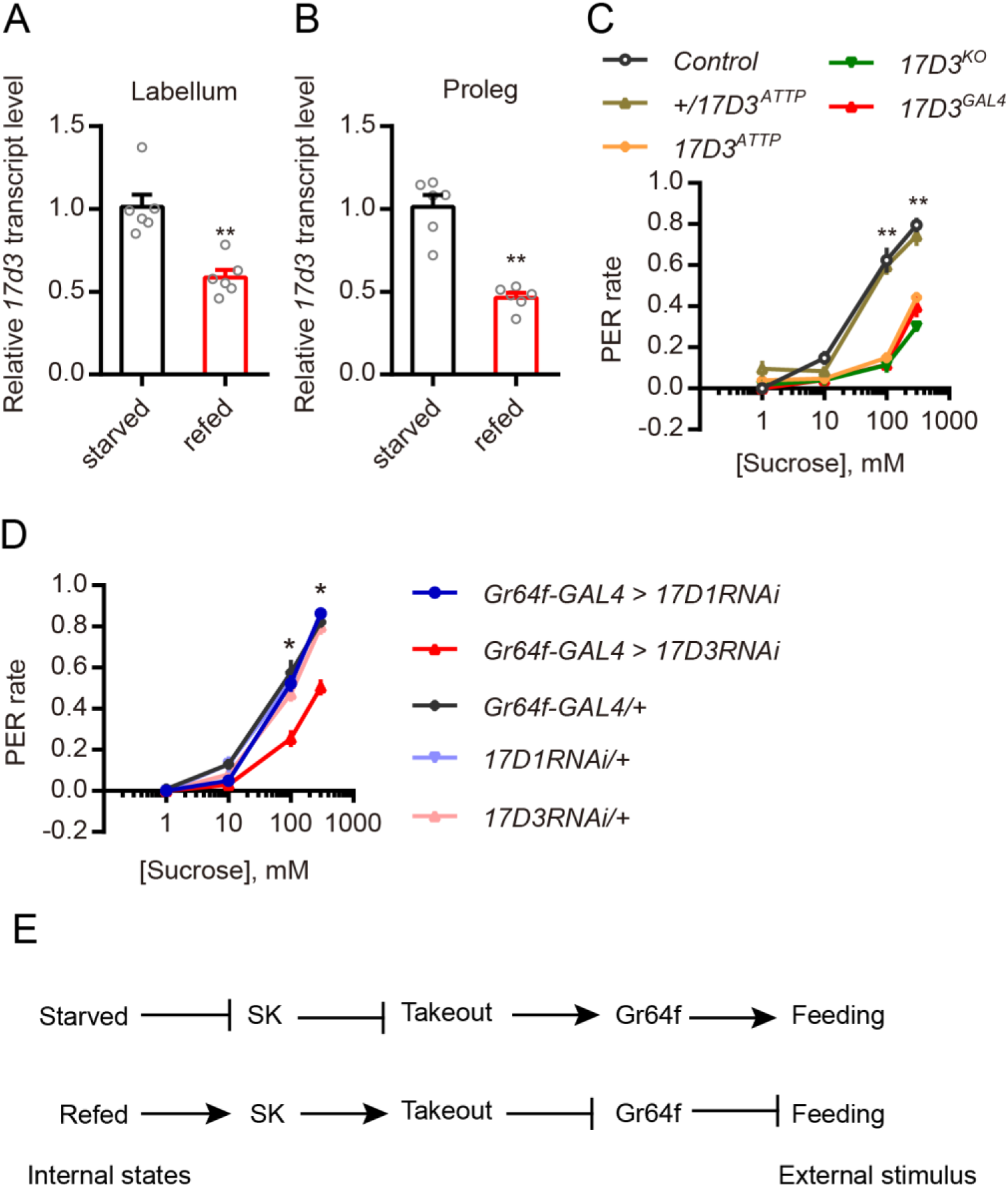
In Drosophila activity of the DSK receptor (CCKLR-17D3) is necessary for PER. (A and B) Refeeding after 24h starvation decreases *17D3* mRNA transcript in the labellum and proleg tarsi. All data are presented as means ± s.e.m. \*\**p* < 0.01; Mann–Whitney test. (C) 17D3 mutants show decreased motivation to feed in PER. \*\**p* < 0.01; Kruskal–Wallis test followed by Dunn’s multiple comparisons test. (D) Silencing of *17D3* gene in *Gr64f-GAL4*-expressing neurons showed decreased motivation to feed in PER. \**p* < 0.05; Kruskal–Wallis test followed by Dunn’s multiple comparisons test. (E) Model of SK as a satiety signal that reflects internal states and inhibits sweet sensation (an external stimulus).

## Discussion

Food seeking and feeding are under complex control by neuronal circuits as well as neuropeptides and peptide hormones ^3, 12, 15, 41, 68,69,71^. Thus, sensory systems, central circuits and interorgan communication, in a nutrient-dependent fashion, contribute to feeding decisions and regulation of food ingestion. In this study, we have analyzed mechanisms of nutrient state-dependent peptidergic regulation of gustatory inputs and feeding.

We show herein that in both the planthopper *N. lugens* and the fly *Drosophila,* SK signaling mediates satiety and decreases sensitivity of gustatory neurons (GRNs) expressing the gustatory sugar receptor Gr64f. Thus, SK release not only decreases food intake ^32, 33, 72^, but also downregulates attraction to sugar. We find that food ingestion diminishes and starvation increases Gr64f expression under control of SK signaling. In the brown planthopper, the gene *takeout* is upregulated by SK, and we showed that knockdown of *takeout* upregulates *Gr64f* and increases feeding in both insects.

We performed additional experiments in *Drosophila* to reveal further SK signaling mechanisms. Calcium and membrane activity in DSK expressing MP neurons in the brain increase after feeding, suggesting that these neurons receive nutrient signals. We could also show that optogenetic activation of DSK neurons in the brain decreases the PER, whereas activation of *Gr64f* expressing GRNs increases the PER. DSK neurons were found to make functional contacts with Gr64f expressing GRNs in the SEZ, and one of the two DSK receptors, CCKLR-17D3, is expressed in Gr64f expressing GRNs in proleg tarsi, labellum and maxillary palps. Furthermore, the *ccklr-17D3* levels are downregulated in these appendages after feeding. In *Drosophila, takeout* knockdown not only increases feeding, but also expression of Gr64f, further suggesting a role of *takeout* in DSK-mediated satiety signaling. Hence, we find that food ingestion activates SK signaling in the two insects and that SK acts to decrease expression of a sweet receptor and thereby diminish food attraction and feeding (Fig. 9E).

Invertebrate SKs and the vertebrate CCKs are known as satiety inducing peptides that regulate food ingestion ^33, 35, 37, 42, 43, 46, 71, 73, 74^. However, these peptides are also known to act in a multitude of other regulatory functions both centrally and in the periphery ^36, 37, 75^. In insects SKs play additional roles in gut motility ^76^, digestive enzyme production and release ^34, 77, 78^, regulation of sexual arousal ^64^, as well as hyperactivity and aggression ^56, 79^. Similarly, in mammals CCK stimulates pancreatic enzyme secretion and release of insulin and glucagon, gallbladder contraction and gut motility, and is implicated in fear, anxiety, and aggression [see ^35, 37, 75, 80^]. In addition CCK has diverse roles in the brain as a neuromodulator regulating other neurotransmitter systems [see ^75^]. Thus, in planthoppers and flies, one can expect that SK acts at several levels and that these actions are coordinated to generate a relevant behavioral and physiological outcome.

In *Drosophila* and planthopper there are cell bodies of SK-producing neurons only in the brain, and to our knowledge no SK peptide is produced in the intestine or other tissues ^64, 81^; see also FlyAtlas2 http://flyatlas.gla.ac.uk, ^82^. We show here that four pairs of posterior DSK interneurons, MP1 and MP3, with wide arborizations in the brain are likely to underlie the regulation of *Gr64f*-expressing GRNs. These MP neurons display increased calcium and spontaneous electric activity after feeding, and we demonstrate here that the MP1 neurons have processes that superimpose GRN axon terminations in the SEZ [see also ^64^]. Furthermore, optogenetic activation of DSK neurons rapidly inhibits the PER suggesting direct neuronal connections. A subset of the brain insulin-producing cells (IPCs) is also known to co-express DSK ^32, 64, 79^, but our experiments exclude these cells in modulation of GRNs. The IPCs may instead release DSK as a circulating hormone to act on the intestine in regulation of digestive enzymes as demonstrated in the planthopper and other insects [see ^34, 77, 78^], but not yet shown in *Drosophila*.

We found that knockdown of *takeout* increases feeding, and also that this upregulates expression of *Gr64f* in planthopper and fly. Interestingly, it has been shown earlier that *takeout* is expressed in GRNs of the labellum in *Drosophila* and that mutant flies are deficient in sugar sensing and regulation of food ingestion ^50^. *Takeout* mutants are also aberrant in their starvation-induced locomotor activity and display increased mortality during starvation ^49, 50^. Furthermore, the gene *takeout* is expressed also in the intestine and fat body, is under control by the circadian clock, and was proposed to link circadian rhythms and feeding behavior ^49^. Since *takeout* encodes a putative juvenile hormone (JH) binding protein, it was suggested that it regulates levels of circulating JH and that this may impart the effects on locomotor activity and food intake, as well as effects on metabolism ^50^. The role of *takeout* in the GRNs in modulation of sugar sensitivity by regulating *Gr64f* in the same cells requires further investigation.

As mentioned, SK acts at several levels of the CNS and periphery and modulates conflicting behaviors such as food search, feeding, sexual arousal and aggression ^32, 56, 64, 79^. These studies show that *Dsk* neurons increase aggression and decrease feeding and mating. The DSK-expressing MP neurons are central in suppression of both feeding (this study), mating ^64^ and aggression ^55^, but aggression was also found dependent on DSK in the IPCs ^56, 79^. Modulation of feeding and mating relies on different downstream circuits. For suppression of male sexual behavior DSK (from MP neurons) acts on male-specific fruitless-expressing P1 neurons that coordinate arousal-related behaviors such as sex, sleep and locomotion ^64^. In parallel, as we show here, the same DSK neurons target GRNs to suppress sweet attraction and inhibit feeding. It is not clear at present how DSK released from IPCs suppresses feeding and what the hormonal DSK targets are ^32^.

Other systems, such as the four widely arborizing SIFamide expressing neurons of the pars Intercerebralis, are also known to coordinate hunger and satiety signals to stimulate appetitive behavior and suppress mating behavior and sleep ^18, 83^. Also this SIFamide system acts at different levels, such as olfactory and gustatory circuits, as well as sleep and activity circuits and fruitless expressing neurons ^18, 84^. Several studies have also shown peptidergic modulation of ORNs and GRNs to promote state-dependent food seeking in *Drosophila*. Thus, nutrient dependent insulin signaling modulates sNPF and TK signaling to alter sensitivity of ORNs ^6, 10, 21^, whereas NPF and sNPF regulate sweet and bitter sensitivity, respectively ^11^. NPF and sNPF are also known to act in other circuits to modulate feeding, metabolism and sleep [see _e. g._ ^24, 25, 26, 27, 29, 30, 31^].

In summary we show that brain DSK neurons, the MPs, by direct inputs regulate sugar sensitivity of DSK receptor-expressing GRNs in response to food ingestion and thereby diminishes food attraction (Fig. 9E). Mechanistically this state-dependent desensitization of specific GRNs is by SK receptor mediated downregulation of Gr64f expression in GRNs, possibly involving action of *takeout*.

## Methods

### Experimental insects and husbandry

The brown planthopper *N. lugens* was reared on ‘Taichung Native 1’ (TN1) rice (*Oryza sativa* L.) seedlings in the laboratory and maintained at 27 ± 1 ^∘^C, with 70 ± 10% relative humidity, under a 16 h: 8 h light dark photoperiod ^85^.

Flies were maintained on standard molasses/cornmeal/yeast/agar food at 25°C on a 12:12 LD cycle with humidity set to 60 ± 5% unless otherwise indicated. The following fly strains were ordered from Bloomington Stock Center: *w^11^*^18^ (Bloomington Stock number: #5905); *Canton-S* (#64349); *Gr64f-GAL4* (#57668; ^86^); *Actin5c-GAL4* (#4414); *UAS-ArcLight* (#51056; ^57^); *UAS-RedStinger* (#8547; ^87^); *UAS-CaLexA* (#66542; ^58^); *UAS-GCaMP7s* (#79032; ^88^); *UAS-Stinger-GFP* (#84277); *UAS-Chrimson, attp18* (#55134; ^59^); *UAS-Chrimson, attp40* (#55135; ^59^); *UAS-NaChBac* (#9467; ^89^); *LexAop-rCD2::RFP, UAS-mCD8::GFP* (#67093); *UAS-nSyb-spGFP*^1–10^*, LexAOP-CD4-spGFP*^11^ (#64314; ^90^); *LexAOP-nSyb-spGFP*^1–10^*, UAS-CD4-spGFP*^11^ (#64315; ^90^); *Δ17D1[Df(1)Exel9051]* (#7762;*17D1^KO^* ^67^). *UAS-DskRNAi* (THU2073), *UAS-17D1RNAi* (THU2656), *UAS-17D3RNAi* (THU2970), and *UAS-Gr64fRNAi* (TH04838.N) were purchased from Tsinghua Fly Center at the Tsinghua University ^91, 92^. *UAS-Kir2.1* ^93^, *LexAop2-IVS-nlstdTomato* ^87^, *Dsk-GAL4, Dsk-LexA, 17D3^KO^, 17D3^GAL4^, Dsk^1^*, and *UAS-mCD8::GFP* have been described previously ^64^. *Dilp2-GAL4* (#37516; ^94^) was kindly provided by Dr. Wei Song. *Gr64f^LexA^* and sugar blind flies (*R1,Gr5a^LexA^;+; Δ61a, Δ64a-f*) were a gift from Dr. Hubert Amrein ^61, 62^. *Dsk^attP^*, *Dsk^LexA(RY)^*, *17D1^attP^*, and *17D3^attP^* were kind gifts from Dr. Yi Rao ^95^.

### Peptide synthesis

Sulfakinin peptides were synthesized by Genscript (Nanjing, China) Co., Ltd. Peptides mass was confirmed by MS and the amount of peptide was quantified by amino acid analysis. The amino acid sequence of the peptides used in this study are: *N. lugens* sulfakinin 1: (NlSK1): SDDYGHMRFamide; sulfakinin 2: (NlSK2): GEADDKFDDYGHMRFamide; sulfated sulfakinin1 (sNlSK): SDDY(SO_3_H)GHMRFamide; sulfated sulfakinin2 (sNlSK2): GEADDKFDDY(SO_3_H)GHMRFamide.

### Gene cloning and sequence analysis

We used the NCBI database with BLAST programs to carry out sequence alignment and analysis. Then we predicted Open Reading Frames (ORFs) with EditSeq. The primers were designed by Primer designing tool – NCBI. Total RNA Extraction was using the TRIzol reagent (Invitrogen, Carlsbad, CA, USA) according to the manufacturer’s instructions. The cDNA template used for cloning was synthesized using the Biotech M-MLV reverse transcription kit and the synthesized cDNA template was stored at −20°C.

The transmembrane segments and topology of proteins were predicted by TMHMM v2.0 (http://www.cbs.dtu.dk/services/TMHMM-2.0/) ^96^. Multiple alignments of the complete amino acid sequences were performed with Clustal Omega (http://www.ebi.ac.uk/Tools/msa/clustalo). Phylogenetic tree was constructed using MEGA 7.0 software with the Maximum Likelihood Method and bootstrapped with 1000 replications ^97^.

### Gene expression profile analysis

For the stage-specific expression study of *Nlsk* and *Nlskr*, total RNAs were extracted from pools of multiple individuals in the following developmental stages: egg, 1^st^ to 5^th^ instar nymphs, BM (brachypterous adult male), MM (macropterous adult male), BF (brachypterous adult female), and MF (macropterous adult female). For the tissue-specific expression study of *Nlskr*, total RNA was isolated from various tissues including head, antenna, wing, proboscis, leg, gut and Malpighian tubule of three days of female adults using TRIzol reagent (Invitrogen).

### Quantitative RT-PCR

The first-strand cDNA was synthesized with HiScript® II Q RT SuperMix for qPCR (+gDNA wiper) kit (Vazyme, Nanjing, China) using an oligo(dT)18 primer and 500 ng total RNA template in a 10 μl reaction, following the instructions. Real-time qPCRs in the various samples used the UltraSYBR Mixture (with ROX) Kit (CWBIO, Beijing, China). The PCR was performed in 20 μl reaction including 4 μl of 10-fold diluted cDNA, 1μl of each primer (10 μM), 10 μl 2 × UltraSYBR Mixture, and 6 μl RNase-free water. The PCR conditions used were as follows: initial incubation at 95°C for 10 min, followed by 40 cycles of 95°C for 10 s and 60°C for 45 s. *N. lugens* 18S rRNA or Drosophila *rp49* were used as an internal control (Supplementary Table S1). Relative quantification was performed via the comparative 2^−△△CT^ method ^98^.

### RNA interference in *N. lugens*

For lab-synthesized dsRNA, *gfp, Nlsk, Nlto* and *NlGr64f* fragments were amplified by PCR using specific primers conjugated with the T7 RNA polymerase promoter (primers listed in Supplementary Table S1). dsRNA was synthesized by the MEGAscript T7 transcription kit (Ambion, Austin, TX, USA) according to the manufacturer’s instructions. Finally, the quality and size of the dsRNA products were verified by 1% agarose gel electrophoresis and the Nanodrop 1000 spectrophotometer and kept at −70 °C until use.

The 4th instar nymph of brown planthoppers was used for injection of 60 nl of 5 μg/μl dsRNA per insect. Injection of an equal volume of *dsgfp* was used as negative control. RNAi efficiency was examined by qPCR using a pool of ten individuals on the 3^rd^ day after dsRNA injections. The insects of the third day after dsRNA injection were used for feeding assay and gene relative expression analysis.

### Feeding assay of *N. lugens*

The animals were food deprived for 5 h before onset of the experiment to ensure that all experimental animals were in the same nutritional state prior to the experiment. This was based on several periods of starvation (2, 5, 12, and 24 h) we tested, from which a starvation for 5 h gave the best results in terms of intraexperiment variation. The animals were starved in the morning and the experiments were done in the afternoon. The feeding assay method and artificial diet was adopted as previously reported with modification ^99^. Briefly, the antifeedant potency of sulfakinins was measured in fourth instar nymphs of *N. lugens*. Prior to injection, the peptides were dissolved in PBS. Individual brown planthoppers were then injected with 40 nl of peptide solution (2.25 pmol/insect) or 40 nl of PBS in the lateral side of the abdomen using a FemtoJet system (Eppendorf-Nethler-Hinz, Hamburg, Germany). Immediately after injection, ten animals were placed in separate plastic containers (9 cm long and 2 cm in diameter), provided with 200 μl of artificial diet and allowed to feed *ad libitum* for 24 hours.

For studying the effect of gene silencing on the feeding behavior of *N. lugens*, the dsRNA-injected 4^th^ instar nymphs were reared on rice seedlings in the laboratory and maintained at 27 ± 1 ^∘^C, with 70 ± 10% relative humidity, under a 16 h: 8 h light dark photoperiod to recover for 2 d. After 5 hours starvation, ten nymphs were transferred into separate plastic containers (9 cm long and 2 cm in diameter), provided with 200 μl of artificial diet and allowed to feed *ad libitum* for 24 hours as mentioned above. The feeding amount of brown planthopper in each feeding chamber was recorded after 24h. The experiment was repeated at least four times.

### RNA-seq analysis

Total RNA of thirty 4^th^ instar nymphs was isolated at day three after *dsNlsk* or *dsgfp* injections in the 4^th^ instar nymphs using a TRIzol reagent (Invitrogen) according to the manufacturer’s protocol. Library construction and sequencing was performed by Novogene with Illumina HiSeq2000 platform (Novogene Bioinformatics Technology Co.Ltd, Beijing, China). Raw sequence data were submitted to the Short Read Archive (SRA) database of NCBI under the accession numbers SRR12460889 (*dsNlsk-1*), SRR12460895 (*dsNlsk-2*), SRR12460894 (*dsNlsk-3*), SRR12460893 (*dsNlsk-4*) and SRR12460896 (*dsgfp-1*), SRR12460892 (*dsgfp-2*), SRR12460891 (*dsgfp-3*), and SRR12460890 (*dsgfp4*).

The raw data were analyzed after filtering the low-quality sequences. Sequences were aligned to the *Nilaparvata lugens* genome (https://www.ncbi.nlm.nih.gov/genome/?term=Nilaparvata+lugens) using Hisat2 v2.0.5. The expression level of genes from the RNA sequencing was normalized by the FPKM method (Fragments Per Kilobase of transcript sequence per Millions base pairs sequenced). This method considers the effect of sequencing depth and gene length for the reads count at the same time and is currently the most commonly used method for estimating gene expression levels. Differential expression analysis was performed using the DESeq2 R package (1.16.1).The clusterProfiler R package was used for Gene Ontology (GO) enrichment analysis and KEGG pathway analysis ^100^.

### Generation of *17D1^GAL4^* knock-in line

To prepare the 17D1^GAL4^ line, we used CRISPR-HDR (clustered regularly interspaced short palindromic repeats – homology directed repair) method based on previous methods ^64^. We chose the upstream and a downstream guide RNAs targeting the part of first exon using the CRISPR Optimal Target Finder: http://tools.flycrispr.molbio.wisc.edu/targetFinder/. In brief, the part of first CCKLR-17D1 coding exon was replaced by GAL4::p65 (Fig. S12A). Firstly, two gRNAs (gRNA1: 5′-GATTTATAAACTCGGGTCGCA-3′; gRNA2: 5′-TCACCGACAGCGGAGATCTC-3′) against CCKLR-17D1 were inserted into pCFD4 as previously described ^101^. We then fused GAL4::p65 into pHD-DsRed (Addgene #51434) between the EcoRI and the NdeI sites. Next, each homologous arm was subcloned into the pHD-DsRed vector too. We injected the modified pCFD4 and pHD-DsRed plasmids into the embryo of *vas-Cas9* flies (# 51324). The correct insertion was confirmed by 3xP3-DsRed screening and recombination accuracy was confirmed by sequencing.

### NSK/DSK antibody

Rabbit anti-DSK antibody was generated by using the peptide N′-FDDYGHMRFC-C′ that corresponds to the predicted DSK-1 and DSK-2 peptides as antigen as previously reported ^64^. We used the DSK antibody to recognize both SK peptides of *N. lugens* and *Drosophila* since the antigen share the same sequence in two species.

### Immunohistochemistry

Unless otherwise stated, fourth instar nymph of brown planthopper and 3-5-day old mated female flies were dissected under phosphate-buffered saline (PBS; pH 7.4) or Schneider’s insect medium (S2) as previously described ^64, 102^. The tissues were fixed in 4% paraformaldehyde in PBS for 30 min at room temperature. After extensive washing with PAT (0.5% Triton X-100, 0.5% bovine serum albumin in PBS), the tissues were incubated in primary antibody for 24 h at 4 °C and in secondary antibody for 24 h at 4 °C. Primary antibodies used: mouse anti-GFP (Sigma-Aldrich Cat# G6539, 1:1000), rabbit anti-RFP (Abcam Cat#ab62341, 1:1000), mouse anti-Bruchpilot (Developmental Studies Hybridoma Bank nc82, 1:30), rabbit anti-DSK (see antibody generation section, 1:100). Secondary antibodies used: donkey anti-mouse IgG conjugated to Alexa 488 (1:500) or Alexa 555 (1:500) and donkey anti-rabbit IgG conjugated to Alexa 488 (1:500) or Alexa 555 (1:500) (Molecular Probes). The samples were mounted in Vectorshield (Vector Laboratory). Images were acquired with Zeiss LSM 700 confocal microscopes, and were processed with Image J software ^103^.

To quantify NlSK level in the brain, we stained the brains with DSK antibody (Fig. 1E and F). The *dsgfp*- or *dsNlsk*-injected samples were processed in parallel and using the same solution and imaged with the same laser power and scanning settings. With the imaged data, we got “Sum Slices” Z-projection of the sub-stacks encompassing whole brain to measure fluorescence intensity (*F_dsgfp_* and *F_dsNlsk_*), then select a small region without signal as the background fluorescence (*B_dsgfp_* and *B_dsgfp_*) using ImageJ. Then we obtained the relative fluorescence of NlSK as the ratio of NlSK signal to the control signal ((*F_dsNlsk_ - B_dsNlsk_*)/(*F_dsgfp_ − B_dsgfp_*)).

### Arclight imaging

Imaging of freshly dissected brain explants of starved (24 hours) or refed (1.5 hours refed after 24 hours starvation) was performed on a Zeiss 710 NLO Axio Examiner confocal microscope using a water immersion objective (Zeiss, Germany). ArcLight was excited with the 488 nm laser. The objective C-mount image was projected onto the 256 × 256 pixel chip controlled by Zen2010 software (Zeiss Germany). Images were recorded at a frame rate of roughly 80 Hz, and depicted optical traces were spatial averages of intensity of all pixels within the region of interest (ROI, in this study, cell bodies of MP1 neurons), with signals processed as previously reported ^57^. Statistical analysis and plotting of the data were performed using Excel and Prism GraphPad.

### CaLexA measurements

Calcium activity of DSK neurons following starvation and refed was measured using the calcium-dependent nuclear import of LexA (CaLexA) reporter system ^58^. Female flies (4–7-days old) carrying Dsk-GAL4 and UAS-CaLexA system were collected. They were then divided in two groups: one group was starved for 24 hours in presence of water and another group was refed 1.5 hours after 24 hours starvation. Following this period, the fly brains were quickly dissected and were processed for immunohistochemistry. The GFP fluorescence were quantified as described above.

### CsChrimson activation

For optogenetic stimulation, tester flies were collected within twelve hours after eclosion and transferred into a vial with regular food containing 200 μM all-trans retinal (116–31–4, Sigma-Aldrich). The vials were covered by aluminum foil to protect from light for 3–5 days before PER test. We immobilized flies on a glass slide with back down so that the proboscis was exposed to the upside and stimulated the animal with red (620 nm, 0.03 mW/mm^2^, Vanch Technology, Shanghai, China) light. Unless otherwise noted, light stimulation was presented continuously throughout the observation period.

Light intensity was measured by placing an optical power meter (PS-310 V2, Gentec, Canada) nearby the location of glass slide.

### Calcium imaging

*In vivo* GCaMP imaging experiments were performed on 4- to 7-day-old female adult flies. GCaMP7s was expressed in DSK neurons under Gr64f-GAL4 control. Flies were starved 24 hours or refed 1.5 hour after 24 hours starvation. The fly brains were rapidly dissected and fixed. GCaMP7s signals from the Gr64f-GAL4 labeled neurons were imaged on Zeiss LSM 710 confocal microscope (Jena, Germany) and processed the images using Image J and Fiji ^104^. For this experiment, we did not monitor in real-time, but processed flies and recorded the cumulative signal. This method was also adopted by previous studies ^105^.

### Extracellular tip recording

7-10 days old females were used for electrophysiological recordings. All flies were kept in dark after eclosion and fed with 200 µM all-trans-retinal for 3 to 5 days. Before the assays, flies were transferred to a tube contained a filter paper with 2ml of all-trams-retinal solution (200 µM all-trans-retinal diluted in 2 ml ultrapure water) for 24 hours, and then some groups were refed for 1.5 hour. Electrophysiological recording from *Drosophila* labellar taste sensilla were implemented as previously reported ^106^. A reference electrode was inserted into dorsal thorax of fly, and proboscis was fully extended. The recording electrode was approached to the tip of a single L-sensilla on labellum and covered ∼ 50% of the total shaft length. Beadle-Ephrussi Ringer solution (B&E) was used as the reference electrode electrolyte, which contained 7.5 g NaCl, 0.35g KCl, and 0.279 g CaCl2·2H2O in one liter of ultrapure water. Using 30 mM tricholine citrate solution (TCC) solved with 100 mM sucrose as the recording electrode electrolyte. Both reference and recording electrodes were capillary glass (borosilicate glass with filament, BF120-69-15; O.D.: 1.2mm, I.D.: 0.69mm) that was pulled on a P-97 puller (Sutter Instrument Corp). The recording electrode was connected to an amplifier (TastePROBE DTP-02, SYNTECH) and a data acquisition device (Axon Digidata 1550B, Molecular Device) under the control of Axon pCLAMP 10.6 software (Molecular Devices). Data was analyzed with Clampfit 10.7 software.

### Proboscis extension response (PER) assays

We performed the PER assays at about 2:00-5:00 pm (ZT6–ZT10) at room temperature. 3-5-day-old females were used for the assays. Before the assays, flies were starved for 24 hours, and refed for 1.5 hour. We first anesthetized flies with carbon dioxide and stuck them on microscope slides. After one hour recovery, we tested flies with water. Proboscis were touched by a water drop, and if the fly did not extend its proboscis in three seconds, we performed the assays with different concentrations of sucrose. Two values were used in the PER assays. A score of 1 means a fly that extended its proboscis and ingested after being fed the sucrose water drop. If not, the score of that fly is 0. We averaged the scores of 5-10 flies as one replicate.

### Capillary feeding (CAFE) assay

This method was modified from Ja et al. ^60^. A vial (9 cm height × 2 cm diameter), filled with 5 ml of 1% agarose to provide water for the flies, was used for this assay. Capillaries (5 μl, VWR International) were exchanged daily with new ones containing fresh food solution (5% sucrose and 5% yeast extract dissolved in water). Typically, 24-h feeding of every fly is shown after 1 or 2 d of habituation in this assay. The amount of consumed food minus evaporation was quantified.

### Statistics

We used the GraphPad Prism 7 software package to generate graphs and statistically analyze data. The Mann Whitney test was used to compare two columns. The Kruskal-Wallis one-way ANOVA test, followed by Dunn’s post test, was used to compare multiple columns of data. All data are presented as mean ± s.e.m. The sample sizes and statistical tests used for each experiment are stated in the figures or figure legends.

## ACKNOWLEDGMENTS.

We thank Yi Rao (Peking University) and Wei Song (Wuhan University) for sharing fly strains; Other members of the Wu laboratory for helpful discussions. This research was supported by the National Natural Science Foundation of China (No. 32022011 & 31772205).

## AUTHOR CONTRIBUTIONS

S.F.W., Y.F.P. and D.R.N. conceived the study and wrote the manuscript. S.F.W., D.G., C.G., Y.F.P., C.X.L., and Y.J..Z performed all experiments and analyzed the data. S.F.W., S.Z., J.L. Y.L.J., C.Y.L., J.Y.M. and C.F.G. performed brown planthopper experiments and analyzed RNA-seq data.

